# Depletion Assisted Hemin Affinity (DAsHA) Proteomics Reveals an Expanded Landscape of Heme Binding Proteins

**DOI:** 10.1101/2022.10.11.511733

**Authors:** Hyojung Kim, Courtney M. Moore, Santi Mestre-Fos, David A. Hanna, Loren Dean Williams, Amit R. Reddi, Matthew P. Torres

## Abstract

Heme *b* (iron protoporphyrin IX) plays important roles in biology as a metallocofactor and signaling molecule. However, the targets of heme signaling and the network of proteins that mediate the exchange of heme from sites of synthesis or uptake to heme dependent or regulated proteins are poorly understood. Herein, we describe a quantitative mass spectrometry-based chemoproteomics strategy to identify exchange labile hemoproteins in human embryonic kidney HEK293 cells that may be relevant to heme signaling and trafficking. The strategy involves depleting endogenous heme with the heme biosynthetic inhibitor succinylacetone (SA), leaving putative heme binding proteins in their *apo-*state, followed by the capture of those proteins using hemin-agarose resin and finally elution and identification by mass spectrometry. By identifying only those proteins that interact with high specificity to hemin-agarose relative to control beaded agarose in a SA-dependent manner, we have expanded the number of proteins and ontologies that may be involved in binding and buffering labile heme or are targets of heme signaling. Notably, these include proteins involved in chromatin remodeling, DNA damage response, RNA splicing, cytoskeletal organization and vesicular trafficking, many of which have been associated with heme through complimentary studies published recently. Taken together, these results provide support for the emerging role for heme in an expanded set of cellular processes from genome integrity to protein trafficking and beyond.

## 1 | INTRODUCTION

Heme *b* (iron protoporphyrin IX), hereafter referred to as heme, is an essential cofactor and signaling molecule^1–9^. As a cofactor, heme facilitates diverse processes spanning electron transfer, chemical catalysis, and gas binding and transport^5^. As a signaling molecule, heme can bind and regulate the activity and/or expression of a handful of transcription factors ^1,10–15^, kinases ^16–19^, cell surface receptors ^20^, and ion channels ^21–23^. Given that heme is also potentially cytotoxic, cells must tightly regulate its concentration and bioavailability ^24–26^. The factors regulating the intracellular concentration of heme are well understood, including the mechanisms and atomic resolution structures of all eight heme biosynthetic enzymes and heme degrading heme oxygenases (HO), HO-1 and HO-2 ^6–9^. However, the molecules and mechanisms that regulate heme bioavailability are comparatively less well understood.

A useful framework for understanding the availability and mobilization of heme in trafficking and signaling is to conceptually partition total cellular heme between exchange inert and labile heme binding sites ^8,27–29^. Inert heme cannot readily exchange and typically associates with high affinity, buried binding sites, *e.g.* in cytochromes and globins. Labile heme (LH) can exchange on time scales relevant for physiological heme trafficking, signaling, and buffering. The speciation of LH, including the proteins that bind it, and the targets of heme signaling to which LH may be mobilized to, are largely unknown.

There are numerous analytical methods to measure total heme in cell and tissue samples, including HPLC, mass spectrometry (MS), UV/vis spectrophotometry, or indirectly via fluorescence spectroscopy of de-metallated heme, protoporphyrin IX ^8,30^. Moreover, identification of high affinity exchange inert hemoproteins can be made on the basis of heme co-purifying with the protein when isolated from native or heterologous sources. However, it is exceedingly challenging to identify proteins that constitute the LH pool and targets of heme signaling using traditional biochemical methods due to facile heme exchange and dissociation upon cell or tissue lysis.

To better understand LH, we and others have developed fluorescence and activity-based heme reporters that can bind exchangeable cellular heme ^27,29,31–35^. When coupled with gene deletion or overexpression screens, or screens for drugs, toxins, or various growth conditions and environmental stressors, these heme sensors can reveal new genes and pathways that control LH and heme bioavailability in cells and subcellular compartments. Indeed, by coupling the heme sensors with various genetic, drug, stress, and toxin screens, we previously identified key proteins, *e.g.* glyceraldehyde phosphate dehydrogenase (GAPDH)^29,36^ and GTPases that regulate mitochondrial dynamics and mitochondrial-ER contact sites ^37^, nucleic acids, *e.g.* ribosomal RNA (rRNA) guanine quadruplexes (G4s)^38^, small molecules, *e.g.* nitric oxide (NO)^29^, and heavy metal stress, *e.g.* lead ^39^, as factors that regulate or alter LH and heme availability.

Previous studies have reported the heme-binding competent portion of the proteome in a variety of organisms, including humans ^40–42^. Most of these studies rely on the enrichment of proteins that bind to hemin-agarose. However, hemin-agarose cannot enrich heme-binding proteins whose heme binding sites are otherwise occupied by heme, thereby reducing sensitivity of hemin affinity-MS strategies. Moreover, it is well established that many proteins interact with hemin-agarose despite not being *bona fide* heme dependent or regulated proteins, or heme homeostatic factors ^9^. Due to these limitations, we hypothesized that the comprehensive hemoproteome has yet to be fully discovered.

Herein, we developed a SILAC (Stable Isotope Labeling with Amino Acids in Cell culture) MS-based proteomics strategy to identify physiologically relevant exchange labile heme binding proteins that could contribute to buffering cellular heme or that could act as sources or targets of heme signaling. To distinguish physiological from non-physiological hemin-agarose binding, we devised a strategy that involves first depleting labile heme with the heme synthesis inhibitor succinylacetone (SA), then identifying proteins that interact specifically with hemin-agarose. The method detects SA-dependent protein interactions with hemin-agarose. In total, the approach reported herein represents the first study of its kind to comprehensively identify the LH proteome in a human cell line without the typical confounding factors associated with heme affinity chromatography.

## 2 | MATERIALS AND METHODS

### SILAC cell culture

Human embryonic kidney HEK293 cells were obtained from the laboratory of Loren D. Williams (Georgia Institute of Technology). HEK293 cells were cultured for 6 cell passages in SILAC media consisting of Powdered DMEM Medium for SILAC (Thermo, #88425), L-leucine (105 mg/L, Sigma, #L8912), L-proline (200 mg/L, Sigma #L5607), 2% penicillin streptomycin (VWR, #97063-708), and 10% Dialyzed Fetal Bovine Serum (R&D Systems, #S12850) supplemented with either 0.798 mM “heavy” L-lysine-13C6, 15N2 (Lys8, Cambridge Isotope Laboratories, #CNLM-291-H), and 0.398 mM “heavy” L-arginine-13C6, 15N4 (Arg10, Cambridge Isotope Laboratories, #CNLM-539-H) for SILAC heavy group or the same concentrations of “light” L-lysine (Lys0, Sigma, #L8662), and L-arginine (Arg0, Sigma, #A6969) for SILAC light group at 37 °C in 5% CO2.

### Succinylacetone (SA) treatment and Cell lysis

One pair of SILAC “light” and “heavy” groups were treated with 500 μM SA for 48 hours (SA+) and the other pair were treated with DI water (SA−) with 3 biological replicates. The SILAC-labeled HEK293 cells with or without SA were harvested in 1X PBS and lysed by 3 cycles of freeze (−80 °C)-thaw method in 10 mM phosphate buffer pH 7.4 with 50 mM NaCl, 5 mM EDTA (G-Biosciences, #015E-F), 1X ProteaseArrestTM (G-Biosciences, #71003-168), 1 mM PMSF (G-Biosciences, #786), and 0.1% Triton X-100 (AMRESCO, #0694).

### Heme Affinity Chromatography (HAC)

To perform HAC, 1.5 mg protein lysate was incubated with 150 μl bead bed of pre-equilibrated hemin-agarose (Sigma, #H6390) for SILAC “heavy” group and the same bead bed volume of pre-equilibrated Sepharose (Sigma, #4B200) in HAC buffer (10 mM phosphate buffer pH 7.4 with 50 mM NaCl, 5 mM EDTA, 1X ProteaseArrestTM, and 1 mM PMSF) for 1 hour at 20 rpm at a rotator at room temperature. The incubated beads were washed 3 times with HAC buffer followed by eluted in 1 M imidazole (Sigma, #IX0005) in HAC buffer.

### Total Heme Measurements

Measurements of total heme were accomplished using a fluorometric assay designed to measure the fluorescence of protoporphyrin IX upon the release of iron from heme, as previously described ^43^. After harvesting HEK293 cells, the cells were counted using an automated TC20 cell counter (BioRad). At least 500,000 cells were resuspended in 400 μl of 20 mM oxalic acid overnight at 4 °C protected from light. After the overnight incubation, an equal volume of 2 M warm oxalic acid was added to give a final oxalic acid concentration of 1M. The oxalic acid cell suspensions were split, with half the cell suspension transferred to a heat block set at 100 °C and heated for 30 min and the other half of the cell suspension kept at room temperature (25 C) for 30 min. All suspensions were centrifuged for 3 min on a table-top microfuge at 21,000g, and the porphyrin fluorescence (ex: 400 nm, em: 620 nm) of 200 μl of each sample was recorded on a Synergy Mx multi-modal plate reader using black Greiner Bio-one flat bottom fluorescence plates. Heme concentrations were calculated from a standard curve prepared by diluting 2.5 to 200 nM hemin chloride stock solutions in 0.1 M NaOH into oxalic acid solutions prepared the same way as for the cell samples. To calculate heme concentration, the fluorescence of the unboiled sample (taken to be the background level of protoporphyrin IX) is subtracted from the fluorescence of the boiled sample (taken to be the free porphyrin generated upon the release of heme iron). The cellular concentration of heme is determined by dividing the moles of heme determined in this fluorescence assay and dividing by the number of cells analyzed, giving moles of heme per cell, and then converting to a cellular concentration by dividing by the volume of a HEK293 cells, assumed to be 1.2 pl ^44^.

### Labile Heme Measurements

Labile heme was measured using a previously reported genetically encoded heme sensor, HS1 ^29,37–39,43,45,46^. HS1 was expressed using the pcDNA3.1 plasmid and driven by the CMV promoter, as previously described ^38,43^. For heme sensor transfections, 1/24 of the HEK293 cells from a confluent T75 flask were used to seed 2 ml cultures in each well of polystyrene-coated sterile 6-well plates (Greiner). Once the cells were 30% to 50% confluent, typically 1 to 2 days after seeding, the media was changed to 2 mls of fresh regular media (DMEM, with 4.5 g/l glucose and without L-glutamine and sodium pyruvate, supplemented with 10% v/v FBS (heat inactivated) 60 min before transfection. For each well to be transfected, 100 μl of a plasmid transfection master mix was added. The master mix was prepared by mixing, in order, 600 μl OptiMEM (100 μl per well up to 600 μl), 2 μg of heme sensor plasmid, a volume of Lipofectamine Plus Reagent equal in μl to the μg of plasmid DNA used, and a volume of Lipofectamine LTX transfection reagent that is double the volume of the Lipofectamine Plus Reagent. Before adding the transfection mixture to each 6-well plate, the master mix was mixed by gentle pipetting and allowed to incubate at room temperature (25 °C) for 5 min. Forty-eight hours after transfection, the media was changed to fresh regular media, regular media with 500 μM succinylacetone (SA), regular media with 350 μM aminolevulinic acid (5-ALA), or heme deficient and SA (HD + SA) treated media. HD media was prepared by using heme depleted FBS, which was produced by incubating it with 10 mM ascorbic acid in a 37 °C in a shaking incubator at 200 RPM for 8 hrs ^16,43^. Loss of heme was monitored by measuring the decrease in Soret band absorbance at 405 nm. Eighty milliliters of ascorbate-treated FBS was then dialyzed against 3 L of PBS, pH 7.4, three times for 24 h each at 4 C, using a 2000 MWCO membrane (Spectra/Por 132625). Dialyzed FBS was then filter sterilized with a 0.2 μm polyethersulfone filter (VWR 28145-501) and syringe (VWR 53548-010). The cells were then cultured for an additional 48 hrs before harvesting for flow cytometric analysis of sensor fluorescence.

For flow cytometric analysis of sensor fluorescence, the cells were washed and resuspended in 1 ml of PBS and transferred to a 1.5 ml microfuge tube and pelleted at 400g and 4 C for 4 min. Supernatant was decanted and the cells were resuspended in 500 μl PBS. The cell suspensions were filtered through a 35 μm nylon filter cap on 12 × 75 mm round bottom tubes (VWR/Falcon 21008-948). Flow cytometric measurements were performed using a BD LSR Fortessa, BD LSR II, or BD FACS Aria Illu flow cytometers equipped with an argon laser (ex 488 nm) and yellow-green laser (ex 561 nm). Enhanced green fluorescent protein was excited using the argon laser and was measured using a 530/30 nm bandpass filter, whereas mKATE2 was excited using the yellow-green laser and was measured using a 610/20 nm bandpass filter. Data evaluation was conducted using FlowJo v10.4.2 software. Single cells were gated by size (FSC versus SSC) and only mKATE2 positive cells that had median mKATE2 fluorescence were selected for ratiometric analysis, which typically corresponded to ~5000 cells.

HS1 sensor heme occupancy was determined according to equation 1 ^29,43^.

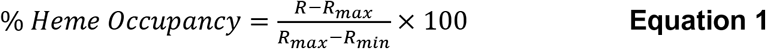

R_min_ and R_max_, the median eGFP/mKATE2 sensor fluorescence ratios when the sensor is depleted of or saturated with heme, respectively, was determined from cultures grown in HD + SA media or media supplemented with 5-ALA. “R” is the median sensor ratio in regular media or regular media supplemented with SA.

### Trypsin digestion

The eluates of SILAC “heavy” and “light” were combined for each SA+ and SA-group respectively followed by reduction in 5 mM TCEP (VWR, #97064-848) for 1 hour and alkylation in 10 mM iodoacetamide (VWR, #97064-926) in dark for 30 minutes. Protein pellet of each SA+ and SA-group was obtained by removing the aqueous and organic layers consecutively following the addition of 4 volumes of methanol (Sigma, #34860), 1 volume of chloroform (ACROS Organics, #423550250), and 3 volumes of water (Sigma, #270733) and centrifugation at 4 °C at 21,000 × g for 10 minutes. The protein pellets were resuspended in 100 mM Tris-HCl pH 8, 1 M ammonium bicarbonate, 6 M urea followed by incubation for 1 hour at 37 °C, and dilution of the urea down to 1.5 M with LC-MS grade water. The resuspended protein samples were enzymatically digested by trypsin with 1:20 (w/w) of enzyme to substrate ratio at 37 °C overnight.

### High-pH RPLC fractionation for whole proteome (WP)

The SILAC-labeled peptides were pre-fractionated by high pH reverse phase liquid chromatography using an Agilent 1100 HPLC system with Waters XBridge C18 reverse phase column (4.6 × 250 mm, 3.5 μM, 130 Å) with 10 mM ammonium formate (solvent A, Fluka, #17843) and 10 mM ammonium formate in 90% acetonitrile (solvent B, EMD Millipore, #AX0145) adjusted to pH 10 using ammonium solution (EMD Millipore, #105426). Fractions were collected every 1 minute at a flow rate of 700 μl/min using a gradient method (from 0% to 75% solvent B in 60 min). The collected fractions were frozen at −80 °C followed by CentriVap lyophilization.

### LC-MS/MS analysis

Peptide fractions were analyzed by LC-MS/MS using a Q Exactive Plus orbitrap mass spectrometer equipped with UltiMate 3000 LC system (Thermo). The lyophilized fractions were resuspended in 0.1% formic acid (FA) in 5% acetonitrile (ACN) and loaded onto NanoViper trap column (75 μm I.D. × 20 mm) packed with AcclaimTM PepMap 100 C18 (3 μm, 100 Å). The samples were resolved through NanoViper analytical column (75 μm I.D. × 150 mm) packed with Acclaim PepMap 100 C18 (2 μm, 100 Å) at a flow rate of 0.3 μl/min with solvent A (0.1% FA in 2% ACN) and solvent B (0.1% FA in 90% ACN) for 150 minutes. Data-dependent tandem mass spectrometry was conducted with the full scans in the range from 200 to 1,800 m/z using an Orbitrap mass analyzer where automatic gain control (AGC) being set at 3e6 (MS1) and 1e5 (MS2), resolution at 70,000 (MS1) and 17,500 (MS2), and Max IT at 100 ms (MS1) and 50 ms (MS2). The top 15 most abundant precursor ions were selected at MS1 with an isolation window of 4 m/z and further fragmented at MS2 by high energy collisional dissociation (HCD) with normalized collision energies (NCE) of 28.

### MS data analysis

MS raw data were searched against UniProt Homo sapiens (Human) proteome (UP000005640) using Proteome Discoverer 2.0 with 10 ppm MS precursor mass tolerance, 0.02 Da MS fragment mass tolerance, and 0.01 false discovery rate. SILAC protein ratios (“heavy” Hemin-agarose / “light” Sepharose for both SA+ and SA-conditions and “heavy” SA+ / “light” SA-for WP) were calculated based on the ratio of integrated peak areas at MS1 for SILAC “heavy” and “light” peptides. Statistical significance of specific hemin-agarose binding of proteins based on the SILAC protein ratio (“heavy” Hemin-agarose / “light” Sepharose) for both SA+ (n=3) and SA-(n=3) were determined respectively using response screening in JMP 16.0. Proteins with either “heavy” or “light” values only were separately processed and reported as +/− infinite change in specific hemin-agarose binding and protein abundance.

### Physical interaction networks and ontology enrichment

Ontology analyses were accomplished using string-db.org version 11.5. Data were filtered for high confidence interactions and clustered using MCL clustering with inflation parameter 3 (default). In cases where there was significant overlap across multiple terms, ontological terms that provided the greatest coverage with minimal overlap between core groups were identified through manual interrogation and highlighted as described in each figure. Aggregate statistics, further analyses and quantitative graphics were generated using JMP Pro version 16 (SAS Inc.).

## 3 | RESULTS

### 3.1 | Depletion Assisted Hemin Affinity (DAsHA) Proteomics

Labile heme protein complexes can readily exchange heme with other molecules on physiologically meaningful time scales. Such complexes represent sites for buffering excess heme and a source of exchangeable heme for heme dependent or regulated proteins. Hemin-agarose based heme affinity chromatography is a commonly used method to enrich potential heme binding proteins (HBPs). However, labile hemoproteins are expected to bind heme at some fractional level reflecting an equilibrium between its heme bound and free states. This fractional saturation reduces the concentration of *apo* HBPs that are competent to bind hemin agarose and prevents the binding of *holo* HBPs that are already complexed with heme. Theoretically, shifting the equilibrium between *holo* and *apo* HBP states should therefore be achievable by depleting cellular heme with succinylacetone (SA), an inhibitor of the second enzyme in the heme biosynthetic pathway, aminolevulinic acid dehydratase (ALAD) (**Figure 1A,B**). We developed a quantitative MS-based hemo-proteomics workflow that capitalizes on this idea in order to reveal new heme binding proteins that have yet to be discovered through traditional hemin agarose enrichment (**Figure 1C**).

**Figure 1.**
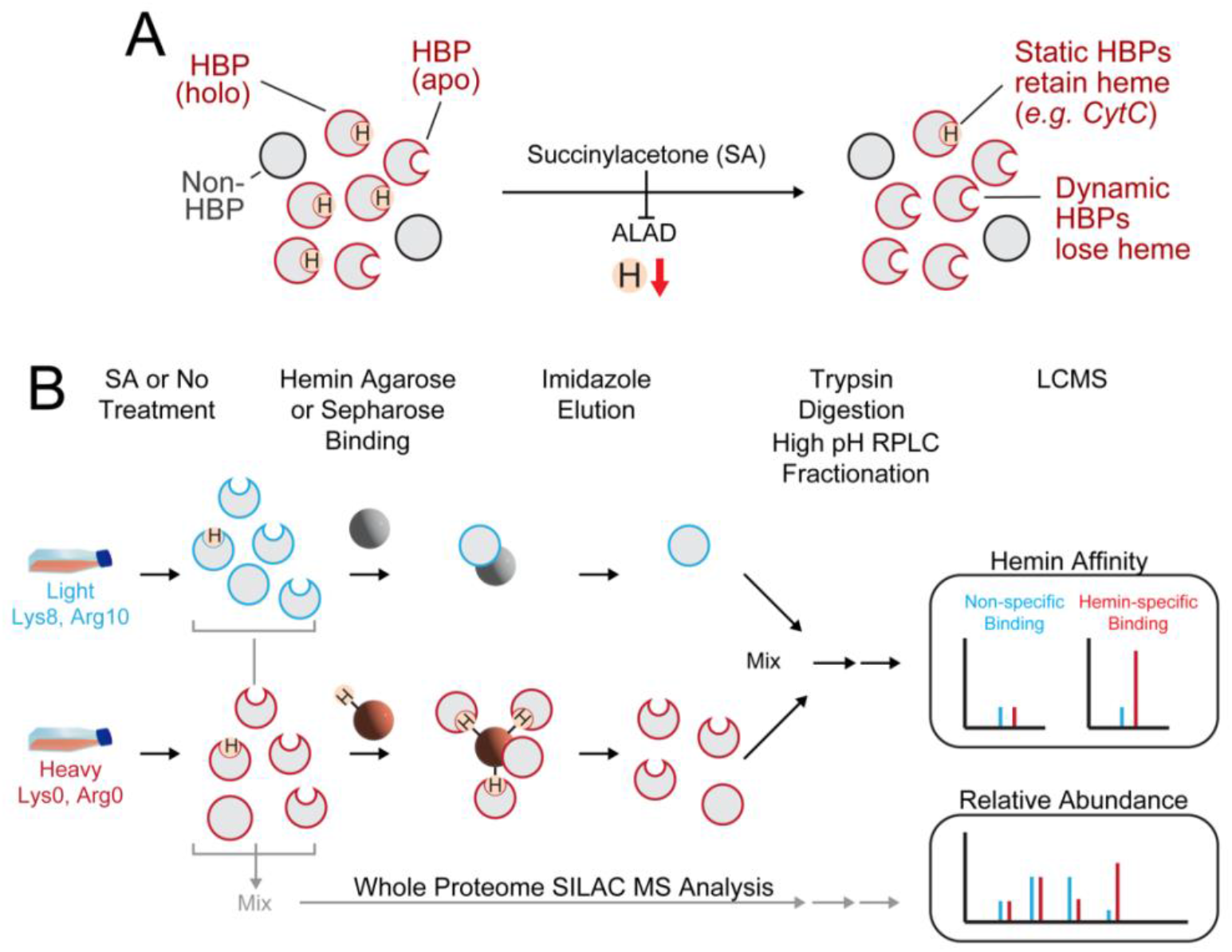
Schematic diagram of DAsHA MS. (A) Succinylacetone (SA) treatment inhibits ALAD to reduce total cellular heme concentrations and increase the frequency of apo-HBPs compared to holo-(heme bound) HBPs. (B) Cells grown in either light or heavy amino acid media are treated with (SA+) or without SA (SA−). The resulting cells are lysed and a fraction is mixed for direct LCMS analysis of proteome wide relative protein abundance (light gray arrows). The remainder of the samples are kept unmixed and exposed in equal amounts to equivalent proportions of either hemin agarose beads (lower path) or beaded Sepharose lacking heme (upper path). After binding and elution with imidazole, the entire eluate from each path is mixed, digested with trypsin protease and fractionated offline followed by LCMS of each fraction. Peptides from proteins binding non-specifically to both bead types have SILAC ratios at or near unity whereas hemin-specific binding proteins exhibit ratios much greater than unity.

In this method, called **D**epletion **As**sisted **H**eme **A**ffinity (DAsHA) MS, cells are grown in the presence or absence of SA, each condition of which is separately grown with light and heavy amino acids (L-lysine and L-Arginine) so as to enable quantitative MS of both affinity enrichment specificity and proteomic analysis of SA+/− cells (*see materials and methods*). In these experiments, light and heavy signals are primarily used to distinguish specific from non-specific binding on hemin agarose versus beaded sepharose (control), respectively. For example, proteins from light or heavy cells conditioned with SA (SA+) are exposed to equivalent amounts of beaded sepharose or hemin agarose, respectively. The resulting eluate from both bead types is then mixed 1:1, digested with trypsin and analyzed by LCMS. Identical processing is carried out in parallel for the SA-condition. The SILAC ratio for each peptide/protein reflects the relative specificity of the detected protein for hemin agarose over control. In parallel, SA+ (heavy) and SA-(light) proteomic samples are also mixed and analyzed *pre-enrichment* by LC-MS/MS to determine changes in protein abundance that may result from SA treatment (*see methods and materials*). The resulting data are overlayed to provide cellular abundance context for each protein detected by affinity purification.

### 3.2 | Heme depletion has a large effect on a small subset of proteins in the human proteome

We carried out DAsHA analysis for three independent biological replicate samples of HEK293 cells (see methods and materials). Heme depletion was accomplished by treating cells with 500 μM SA for 48 hours, which resulted in a 3-fold depletion of total heme (**Figure 2A**) and complete ablation of exchange labile heme as measured by genetically encoded fluorescent heme sensor, HS1 (**Figure 2B**) ^29^. The eGFP to mKATE2 fluorescence ratio of HS1 is a reporter of labile heme, with a high ratio indicating low heme and a low ratio indicating high heme. Ratios are calibrated against cells grown under heme deficient conditions (HD+SA) or heme saturating conditions (media supplemented with the heme biosynthetic intermediate aminolevulinic acid (5-ALA)) (see materials and methods). To establish the effects of this treatment on proteome-wide protein abundance, we carried out whole proteome SILAC MS analysis of each sample. At scale, we found that SA treatment has a relatively minor effect on protein abundance across the entire proteome, with ~93% of the data falling below a 2-fold change in relative abundance between the two conditions (**Figure 2C, Table S1**). This trend was consistent for the established network of known heme-associated proteins within the human proteome, including cellular activities coupled to heme regulation and toxicity and proteins involved in cellular respiration or antioxidant activity (**Figure 2D,E**).

**Figure 2.**
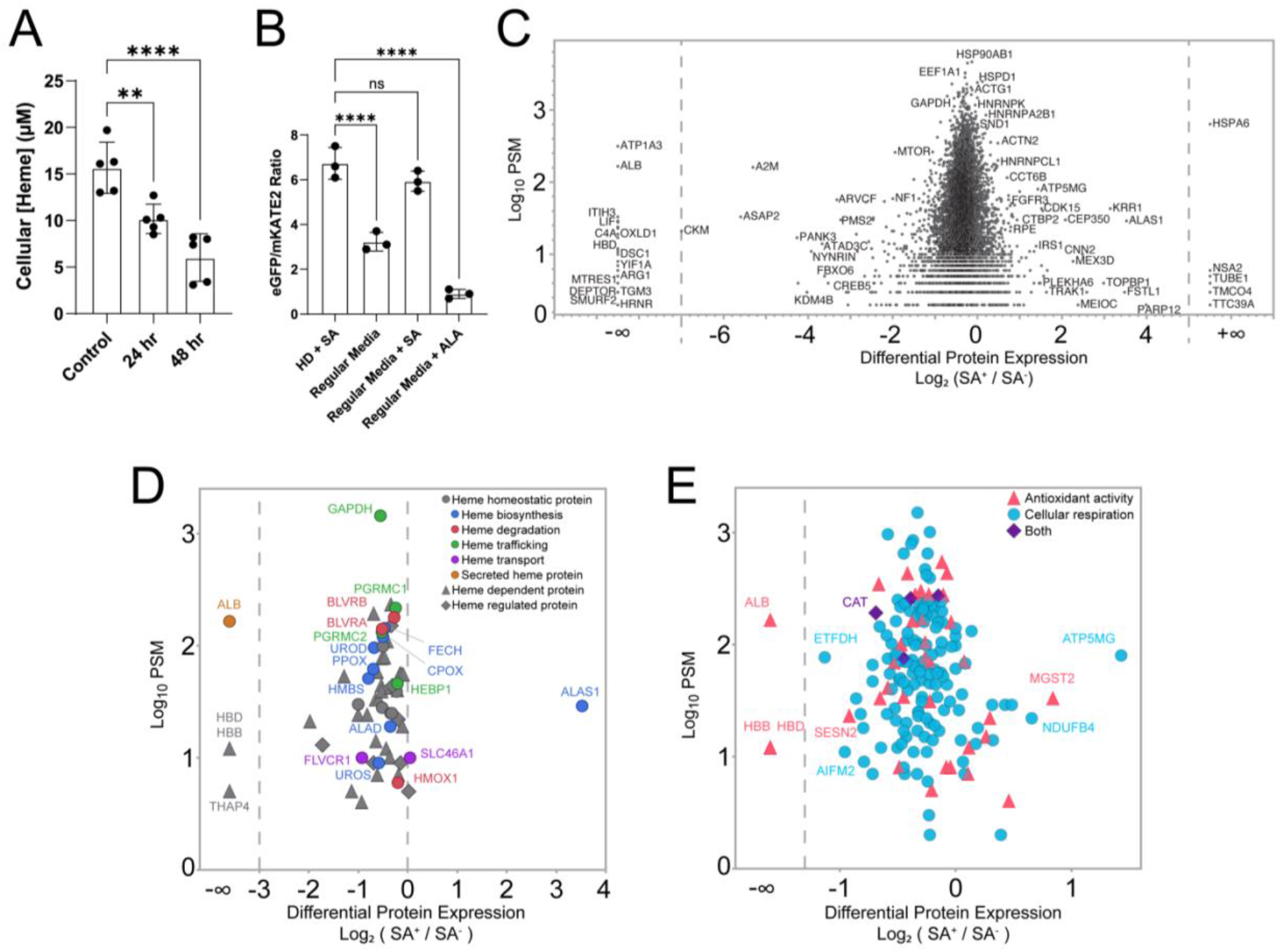
Heme depletion has a minor effect on most, but not all proteins in the proteome. (A) Effect of 24 or 48 hour succinylacetone (SA) treatment of HEK293 cells on total intracellular heme concentrations. (B) Effect of 48 hour SA treatment of HEK293 cells on labile heme as measured by the genetically encoded heme sensor HS1. (C) SILAC MS results for whole proteome measurement of protein differential expression in 48hr SA+ versus SA-conditions. Peptide spectral matches (PSM) correspond with the sum across all peptides for each protein. Proteins visible in only one of two conditions are shown as +/− infinity on the x-axis. (D,E) Subset of proteins from B enriched in specific GO terms associated with cellular heme. Plots are broken into two parts to reduce overcrowding. (See also Tables S1 & S2) Heme homeostatic proteins are defined as those with known effects on heme synthesis, degradation, transport, and trafficking. Heme dependent proteins are those defined as requiring heme for activity. Heme regulated proteins are those that exhibit activities that are modulated by heme but that do not require heme for function.

A small proportion of proteins undergo extreme changes in abundance in response to SA treatment as indicated by an extremely large SILAC ratio or the lack of signal from one of the two samples (see +/−∞) (**Figure 2C-E & Table S2**). We found that in several cases, these proteins either rely on heme binding for stability, such as hemoglobin [HBB] or albumin [ALB] ^47^. In contrast, we also observed dramatic increases in a small subset of proteins, including the heme biosynthetic enzyme Alas1 (5’-aminolevulinate synthase), which undergoes repressed transcription and accelerated protein degradation in response to heme binding and should therefore increase in response to heme depletion (**Figure 2B**) ^48^. Lastly, we also observed extreme abundance changes in several other proteins not previously known to be heme regulated, which may represent a putative set of heme regulated proteins, albeit not necessarily through direct interaction with heme (**Table S3**). These include nuclear, cytoplasmic, cell membrane, golgi, ER, and secreted proteins that may constitute a protein network whose abundance regulation, either through expression or degradation, relies heavily on cellular heme concentrations.

Proteins undergoing extreme abundance changes in response to heme depletion were also highly enriched in phosphoproteins, some of which are linked to immune response, cell export, and endocytic vesicles (**Figure S1**). Together, these data indicate that SA treatment effectively depletes cellular heme, that such depletion produces expected effects on proteins known to be regulated by heme, but has an overall minor effect on the proteome at large. Furthermore, these data uncover a network of phosphoproteins potentially regulated by dynamic cellular heme concentrations.

### 3.3 | Heme depletion boosts the detection of hemin agarose-binding proteins from the human proteome

In parallel with the whole proteome analysis, samples were subjected to hemin agarose enrichment to reveal heme-specific binding proteins wherein specificity is quantified by the ratio of protein binding to hemin agarose versus control sepharose beads lacking conjugated heme. We observed a total of 343 and 480 proteins enriched >2-fold on hemin agarose compared to sepharose in the SA- and SA+ conditions, respectively (**Figure 3A,B & Table S4**). Overall, non-specific binding to sepharose was low, which we confirmed by SDS-PAGE and Coomassie protein staining (**Figure 3C**). After filtering protein IDs by significance (-Log_10_ p-value > 1.3) those proteins identified exclusively upon enrichment with hemin agarose but not in the sepharose controls were classified as *high* hemin specificity while proteins identified in both hemin agarose and sepharose controls (1 < Log_2_ < 10) were classified as *moderate* hemin specificity (**Table S4**). Importantly, despite showing some tendency to bind non-specifically to sepharose, several proteins classified as having moderate specificity were in fact enriched by as much as 70-fold by hemin agarose compared to sepharose.

**Figure 3.**
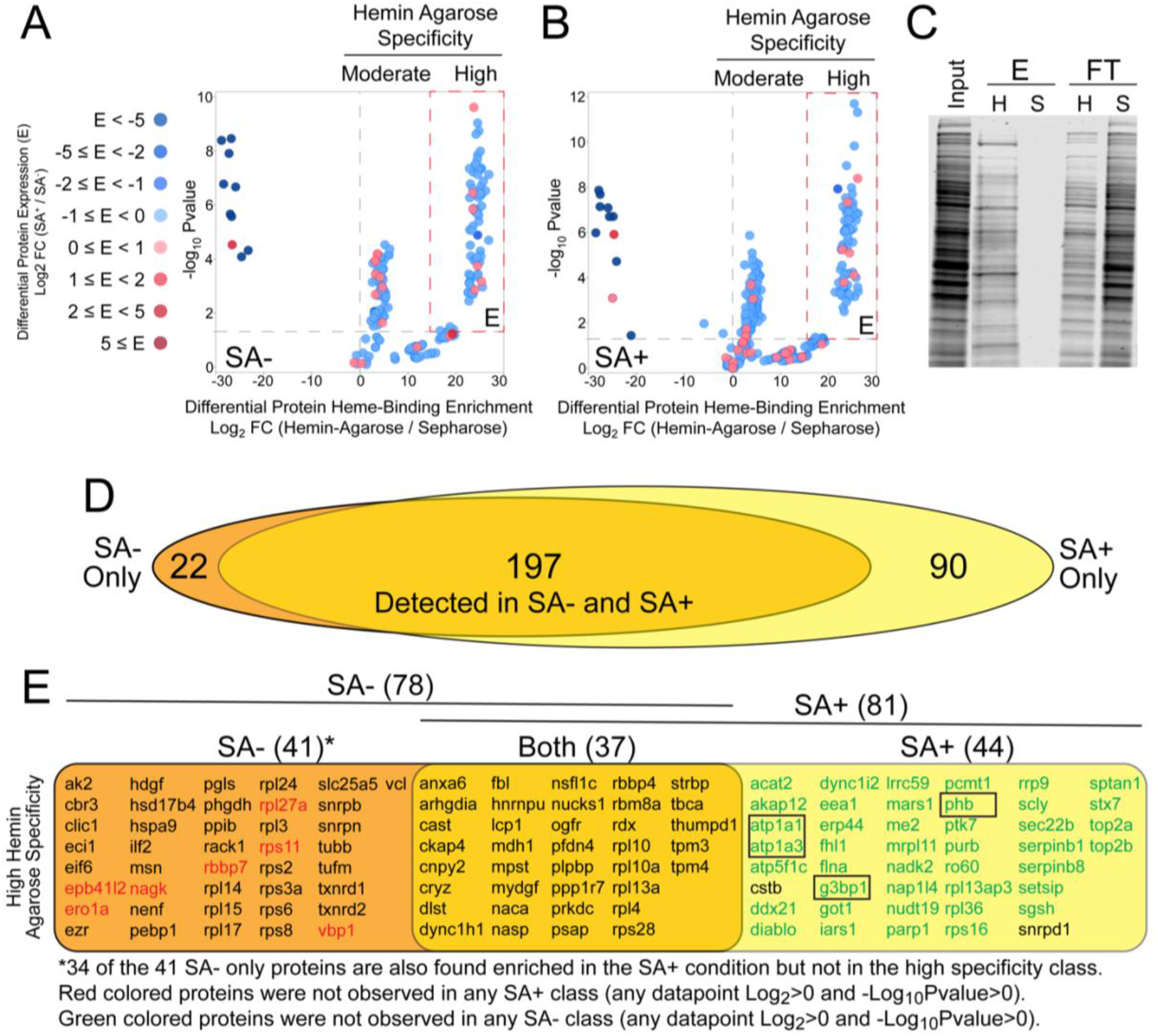
Heme depletion increases the detection of labile heme binding proteins. (A) Plot of hemin agarose-specific binding proteins in the SA-condition. The Log2 SILAC ratio for each protein in the hemin agarose versus Sepharose control. Proteins are clustered into “moderate” and “high” specificity classes where high specificity proteins are those that were detected only in the presence of hemin agarose. Moderate specificity proteins are those that were to varying degrees in both hemin agarose and sepharose control. Note that moderate specificity ranges from log_2_ ~1 (2-fold) to ~6 (~70-fold) enrichment on hemin agarose versus sepharose. Color corresponds with the observed fold change in abundance between SA+ versus SA-conditions (from figure 2 data). Proteins highly enriched on Sepharose over hemin agarose (log_2_ ratios well below 0) are shown as darker colors to denote their general up or down regulation in SA+ versus SA-. Significance determined from 3 independent biological replicates. (C) Coomassie-stained SDS-PAGE showing gross enrichment of proteins on hemin agarose versus Sepharose controls. (D) Venn diagram showing the overlap between proteins detected in the moderate and high hemin specificity classes in SA- and SA+ conditions regardless of significance. (E) List of proteins with high hemin agarose specificity that overlap between SA- and SA+ conditions. (black) proteins observed in either condition when comparing the high specificity class to all classes from the opposite condition; high specificity proteins observed exclusively in the SA-condition (red) or the SA+ condition (green) when comparing to any class (regardless of p-value) in the opposite condition. These therefore represent proteins uniquely detected in either condition. Boxed proteins were validated by immunoblotting (see section 3.4). (See also Table S4)

To evaluate the effect of heme depletion on protein capture, we first compared the comprehensive list of proteins identified (Log_2_ > 1) in the SA-versus SA+ conditions, regardless of significance (309 proteins). While the majority of proteins observed between the two conditions overlapped (197 proteins; 64%), we found a large increase in the number of unique proteins identified only after SA treatment (90 proteins; 29%) in contrast to 22 proteins (7%) observed exclusively in the absence of SA treatment (**Figure 3D**). When restricting the comparison to high hemin specificity proteins we observed 122 proteins in total (SA- & SA+), wherein there was a 30% overlap (37 proteins) between the two conditions and 44 high specificity proteins (36%) in SA+ compared to 41 proteins (34%) in the SA-condition (**Figure 3E**). 42 (95%) of the 44 SA+ high specificity proteins were not enriched at all in the SA-condition, regardless of significance and fold change thresholds. In contrast, all but 7 proteins in the SA-high specificity class were also detected in the SA+ condition regardless of these same thresholds. Taken together, these data demonstrate that depletion of cellular heme prior to hemin agarose enrichment has a bulk advantage in boosting the detection of HBPs in the human proteome. Furthermore, these data support the hypothesis that a subset of the human proteome is involved in interacting (directly or indirectly) with the dynamic labile heme pool in cells.

### 3.4 | Validation of MS-identified heme binding proteins visible only after heme depletion

To validate our MS-based results, we conducted immunoblot analysis of proteins from SA- or SA+ conditioned cells. Samples were again enriched on either sepharose or hemin agarose beads followed by immunoblotting with antibodies to ATP1A-1/2/3, G3BP1, and PHB. GAPDH, which we and others have previously shown interacts with and traffics labile heme ^29,36,49–52^, was included as a positive control. For each of the three test proteins, strong enrichment was observed with hemin agarose over sepharose, consistent with their high specificity classification by SILAC MS (**Figure 4**). Also consistent was our finding that GAPDH interacts mildly with sepharose, which we also observed by SILAC MS (**Table S4**). Each protein, including the GAPDH control, showed an increase in hemin agarose binding in the SA+ condition, which supports the conclusion that heme depletion has a bulk positive effect on boosting enrichment of labile heme binding proteins. These data provided strong support for the validity of the MS results and suggest that the method captures labile heme binding proteins that may not be detected through traditional approaches that do not deplete cellular heme.

**Figure 4.**
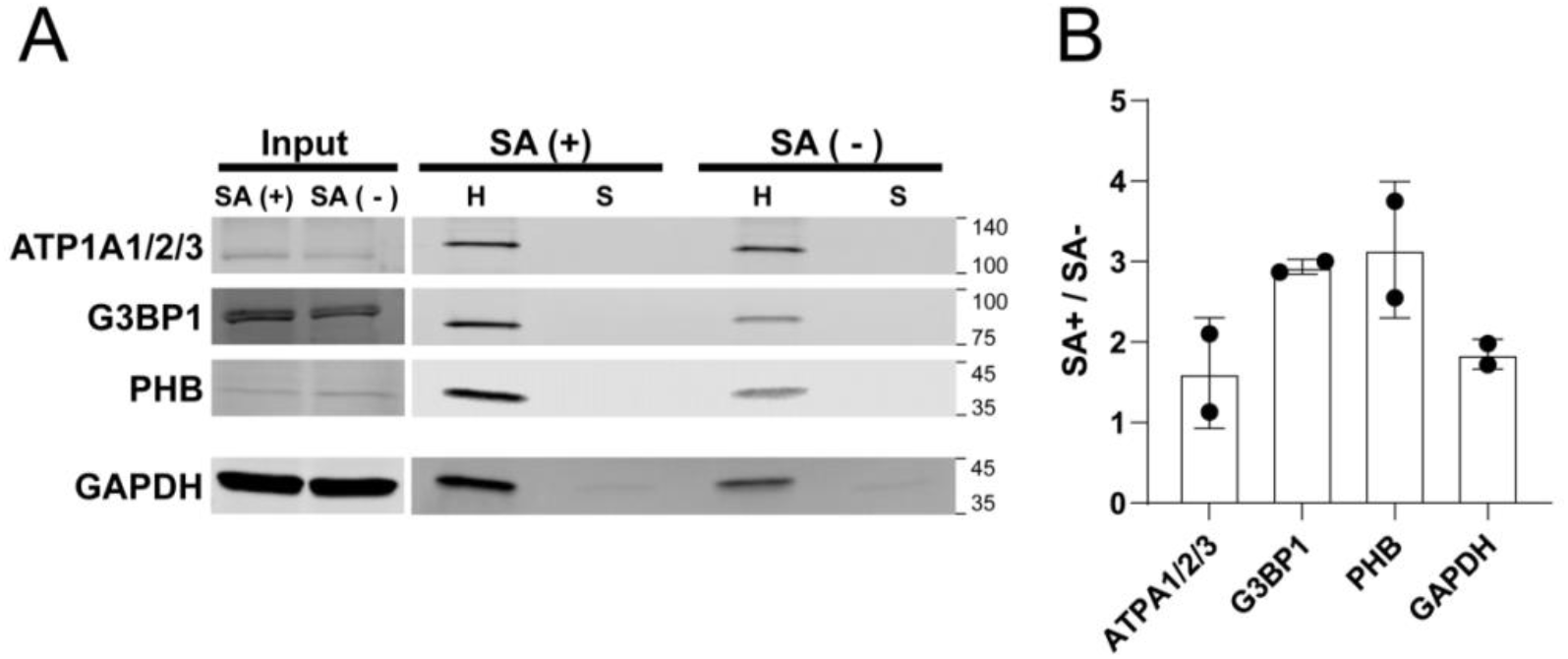
Immunoblot validation of select proteins identified by DAsHA MS. A selected set of proteins identified as hemin agarose specific exclusively in the SA+ condition (ATP1A1/2/3; G3BP1; PHB) were validated by immunoblotting with protein-specific antibodies to confirm their hemin agarose specificity. GAPDH, which is an established heme binding protein was evaluated in parallel as a control. Shown here are the input and elution fractions after enrichment with hemin agarose (H) versus sepharose (S). Note that the tendency for GAPDH to bind sepharose, visible as a band in the sepharose lanes, is also observed in the MS data (see Table S4).

### 3.5 | Ontology enrichment reveals higher than average physical connectivity for heme binding proteins

To gain insight into the human proteomic targets of heme, we conducted ontological enrichment of the comprehensive list of proteins (moderate or high specificity) observed in both SA- and SA+ conditions with significance (p<0.05). Of this list of 218 proteins, 213 matched to a protein in the STRING ontology database. Generally speaking, these proteins show significant enrichment across a wide variety of locations including extracellular space, cytosol, mitochondrion, endoplasmic reticulum, and nucleus, suggesting that protein enrichment was not biased towards one cellular space versus another (**Figure S2**). Strikingly, nearly half of the 213 proteins (~48%) were classified as RNA binding (**Figure S3**). Gene ontology and KEGG pathway enrichment analysis further revealed a broad list of highly overlapping terms with false discovery rates well below the significance threshold (Q<0.05) and with greater than 5-fold enrichment (**Figure 5, Table S5**). We found that a vast majority of ontologies could be explained by physical interactions in protein complexes. For example, biological processes such as *SRP-dependent co-translational protein targeting to membrane*, *nuclear-transcribed mRNA catabolic process*, *viral gene expression*, and *translation initiation* (**Figure 5A**); as well as cell components such as the *polysome* are linked to the ribosome core complex (**Figure 5B**). Similarly, the highly enriched molecular function *threonine-type endopeptidase activity (GO:004298)*, is linked to the 26s proteasome (**Figure 5C**). In light of these and other observations we evaluated the mean node connectivity of each term, focusing on high confidence protein interactions, which revealed a higher than average expected physical connectivity (i.e. *Node Degree in STRING*) for each term type and an overabundance of edges in the physical network across all 213 proteins (805 observed versus 352 expected edges; PPI enrichment p-value < 1E-16) (**Figure 5D, Table S5**).

**Figure 5.**
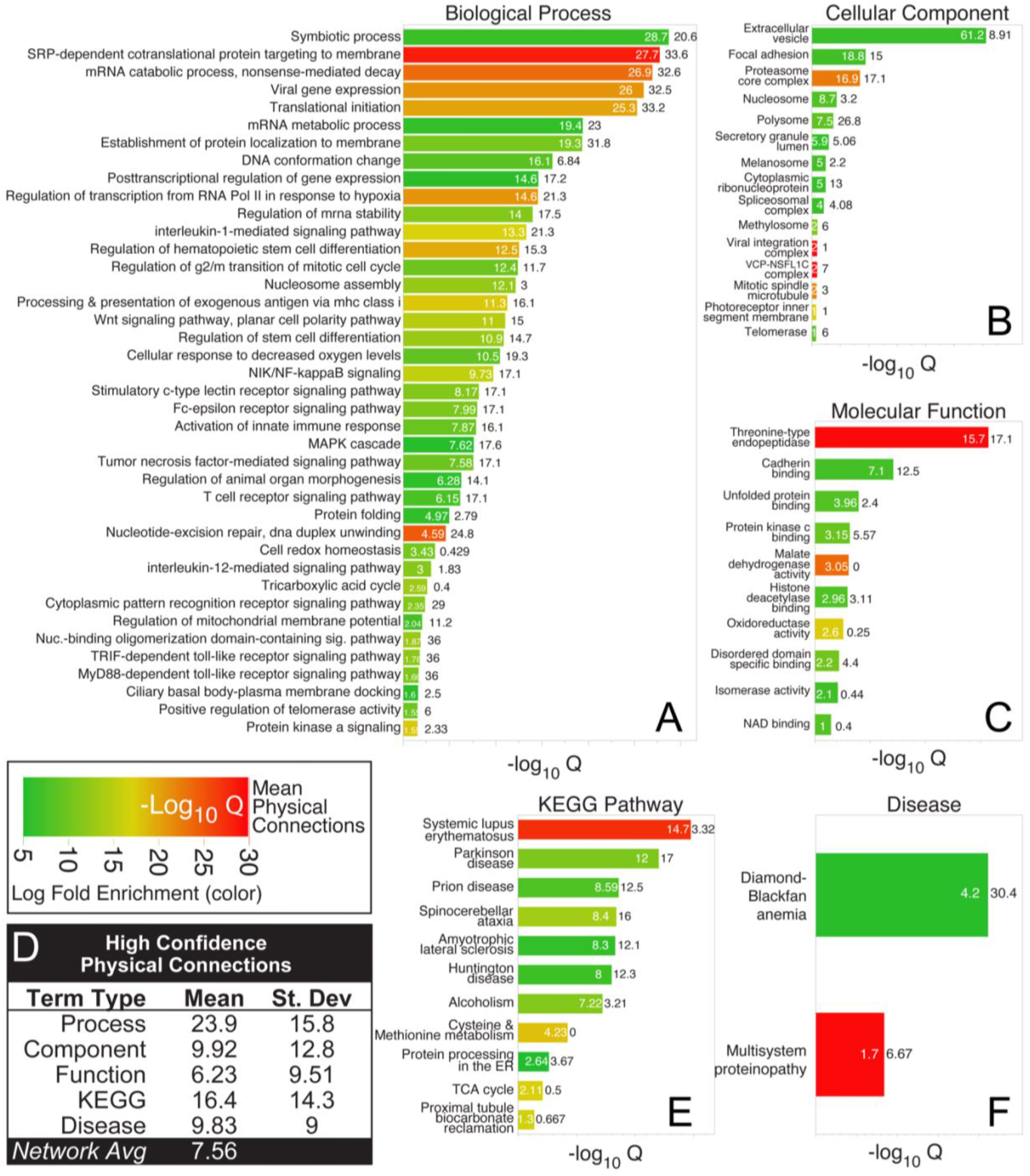
Ontological term enrichment of hemin specific binding proteins. Gene ontology term enrichment for all moderate and high specificity proteins in SA- and SA+ conditions that were enriched above the significance threshold (-Log10 p-value > 1.3): (A) Biological Process; (B) Cellular Component; (C) Molecular Function; (E) KEGG Pathway; and (F) Disease. Each graph shows a bar with embedded value (white numbers corresponding to X-axis value) corresponding to the false discovery rate for the given ontological term. Black values outside each bar correspond to the mean physical connections observed for proteins within each term (corresponding to node degree in STRING). Bar color indicates log fold enrichment (referred to as enrichment “strength” in STRING). (D) Table of high confidence physical connections (min interaction score from STRING = 0.7) observed within each term type shown in A,B,C,E and F. The average node degree for the entire network of high hemin agarose specificity proteins (SA or SA+; 213 proteins) is shown for comparison. (See also Tables S5)

We also observed significant enrichment of multiple KEGG pathways that harbor proteins detected by DAsHA. Most prominent of these was *systemic lupus erythematosus (KEGG ID: hsa05322)*, an ontological pathway linked to the presently untreatable autoimmune disease of the same name which includes nucleosomal histone proteins as well as specific spliceosome components that were detected by DAsHA (**Figure 5E**). Notably, both spliceosome proteins Snrpd1 and Snrpd3 undergo a significant increase in cellular abundance upon heme depletion with SA; and furthermore, Snrpd1 was detected with greater hemin agarose specificity and greater significance in the SA+ condition, providing strong evidence for a role of heme in splicing and in diseases linked to splicing. We also observed significant enrichment in proteins associated with protein misfolding, including Parkinson’s disease (KEGG ID: H00057), Prion disease (KEGG ID: H00061), and Amyotrophic lateral sclerosis (KEGG ID: H00058), within which we routinely observed mitochondrial ATP synthases Atp5f1b and Atp5f1c, among other proteins in the high hemin specificity class (**Table S5**). Similarly, we observed strong enrichment of multi-system proteinopathies (DOID:070355) in the *DISEASES* database, including heterogenous nuclear ribonucleoproteins HNRNPA1 and HNRNPA2B1 and the transitional ATPase VCP (**Figure 5F**).

### 3.6 | Several functional protein complexes revealed by DAsHA

Considering the high degree of protein-protein interaction in our dataset, we next explored whether functional protein complexes may be enriched by DAsHA. We found that 130 of the 213 STRING-mapped proteins (61%) participate in high confidence physical protein interactions (**Figure 6A**). Several distinctive clusters were enriched significantly within this dataset, which we further encapsulated with other proteins of similar function (**Figure 6B**, **Table S6**; corresponding to color-coded clouds in panel A). Ribosomal and proteasomal complexes that have been established as heme interacting previously were the most prominently enriched functional complexes (**Figure 6B**). Also prominent were nucleosomal histones H2B and H4 as well as nucleosome remodeling proteins and the spliceosome. Notably, H2B as well as multiple large ribosomal subunits and spliceosome proteins each exhibit a drastic increase in abundance upon heme depletion (Log_2_ SA+/SA− > 5), suggesting a possible relationship between heme binding and abundance regulation for these proteins (**Figure S4**). Significant enrichment was also noted for additional protein clusters involved in chaperone-mediated autophagy and protein folding, cytoskeleton and vesicular trafficking, and amino acid metabolism (**Figure 6A**).

**Figure 6.**
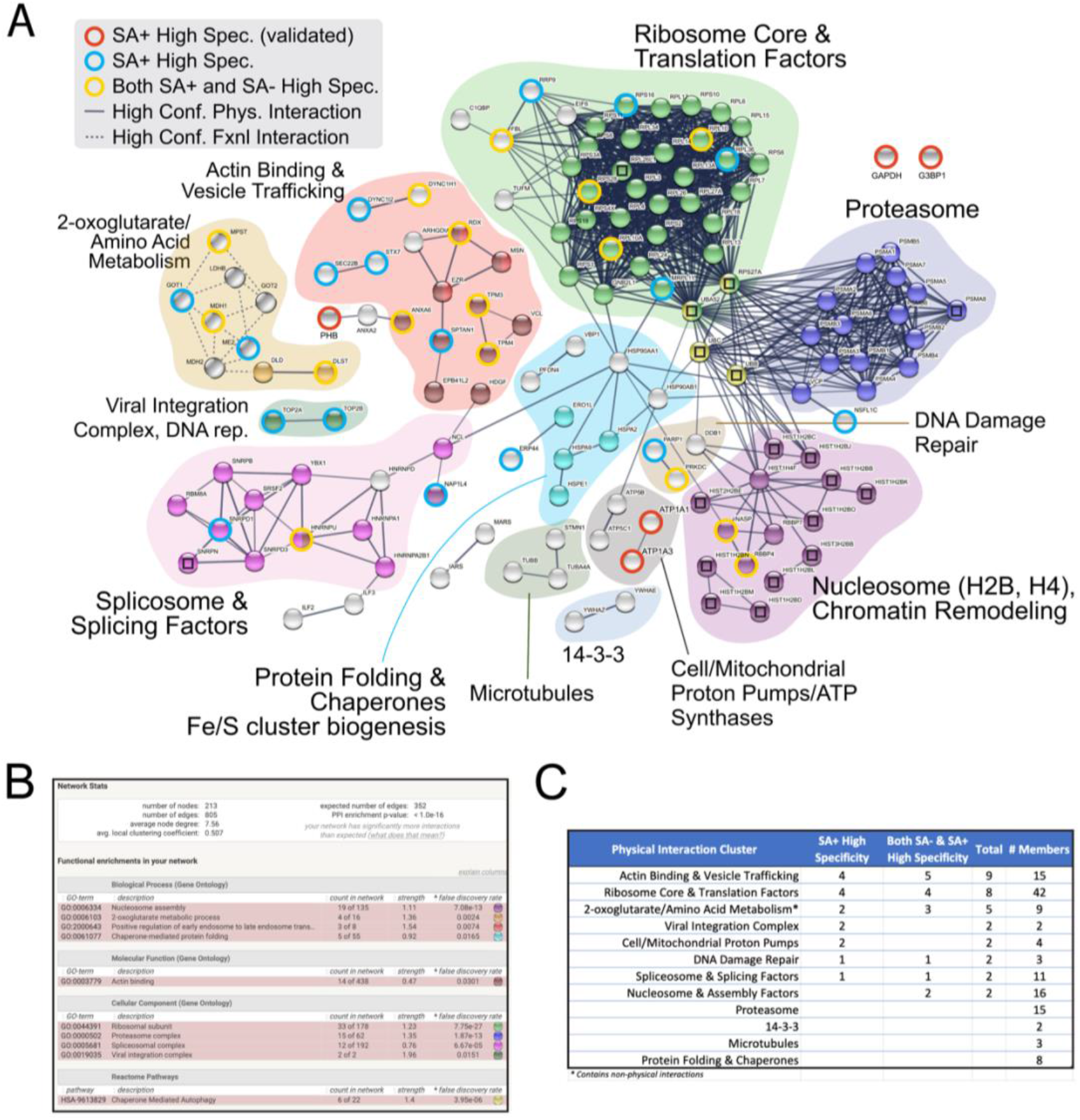
Functional protein complexes identified by DAsHA MS. (A) Network of high confidence physical interactions between moderate or high hemin agarose specific proteins observed in SA- or SA+ conditions (213 proteins; min interaction score from STRING = 0.7) (See also Table S5). Proteins were clustered using the MCL inflation method in STRING and color-coded clouds were manually drawn to encapsulate notable clusters that also include functionally similar proteins not automatically grouped by ontology clustering. Spheres with embedded squares indicate proteins whose detectable peptides were shared with other protein identities (e.g. isoforms). Spheres encircled with red, blue, or yellow rings were detected in the high specificity class (exclusively detected from hemin agarose but not sepharose enrichment) while spheres without rings were detected in the moderate specificity class (2 to 70-fold hemin agarose versus Sepharose enrichment). Not shown are most proteins unconnected by a high confidence physical interaction unless they were explicitly validated (e.g. GAPDH and G3BP1). (B) Snapshot of the STRING network statistics also showing prominent ontologies that were explicitly enriched in the dataset of 213 proteins. Colors correspond to sphere colors in panel A. (C) Table showing the sum of high specificity proteins identified within each ontological cloud.

In most cases we observed that HBPs identified via heme depletion alone (SA+) were found in clusters that were also identified without SA treatment. For example, in clusters associated with actin binding/vesicle trafficking, ribosome, spliceosome, and DNA damage repair, we detected both proteins that require SA for hemin-agarose binding and others that could be detected regardless of SA treatment (**Figure 6C**). In other cases, ontology terms were enriched only after SA treatment, such as the topoisomerases Top2a and Top2b involved in DNA replication and viral integration as well as the sodium/potassium proton pumps Atp1a1 and Atp1a3). These data further convey the expansion of enriched gene ontologies afforded by DAsHA.

Many proteins enriched by hemin agarose (39%) are not classified as having high confidence physical interactions in the STRING database, and in fact we found a slight enrichment (1.3-fold) of SA+ specific proteins in this category (compare 41% to 31% in the set of 213 proteins). Most of these proteins could be broadly linked to general ontologies including: Focal adhesion, mitochondrion/mitochondrial matrix, extracellular exosome, and metabolism where within high confidence functional interactions were noted for 2-oxoglutarate and amino acid metabolism ontologies that exhibited the greatest number of high confidence functional interactions (**Figure S5**).

## 4| DISCUSSION

We have conducted a comprehensive analysis of the heme interactome in human embryonic kidney cells and have expanded the ontological landscape that we understand as the *hemoproteome*. We have revealed a wide range of new proteins and functional protein complexes that associate with heme and may function to buffer toxic heme levels, facilitate heme trafficking, serve as heme-dependent or heme-regulated proteins, or other currently unknown functions. Several of the identified proteins and functional complexes are consistent with recent evidence of heme’s emerging roles in genome, transcriptome, and proteome regulation.

We employed a unique strategy, DAsHA, to improve enrichment of fractionally-saturated HBPs that are otherwise difficult to enrich through traditional hemin agarose chromatography. Coupled with SILAC quantitative mass spectrometry, we significantly improved the capture and detection of hemin agarose-specific binding proteins. The method, by design, did not capture known high affinity heme proteins and enzymes since our heme depletion strategy had more severe consequences on labile heme compared to total heme (**Figure 2A & 2B**; **Table S2**). Rather, the method reveals what we consider to be proteins that interact with dynamic labile heme.

### 4.1 | An emerging direct role for heme in chromatin regulation, DNA replication, and DNA repair

Heme has roles in controlling the expression of specific genes through either direct negative or direct positive transcriptional control mechanisms. In budding yeast, heme binds and activates the transcription factor Hap1, which promotes the expression of genes required for aerobic metabolism and represses hypoxic genes in response to oxygen ^11,14,15,53^. Additionally, heme binding to Gis1, a yeast transcription factor and demethylase orthologous to the mammalian JMJD2/KDM4 family of demethylases, regulates genes involved in the oxidative stress response and carbon metabolism via post diauxic shift (PDS) elements in the promoters of numerous genes activities ^13^. In humans, heme can promote positive or negative regulation of transcriptional repressors and activators such as Bach1 ^13^, p53 ^54^, and Reverbα/β^12,55–57^. Heme binding to transcriptional repressor Bach1 or tumor suppressor protein p53, for example, promotes their proteasomal degradation ^10,13,58^. More recently, it has been shown that heme controls genome function in a parallel mechanism mediated through chromatin remodeling, wherein DNA accessibility at promoter regions at specific locations across the genome (measured by a transpose accessibility sequencing assay) is altered under low heme conditions. resulting in changes in transcription ^59^. The mechanism underlying this heme dependent chromatin remodeling is presently unknown. Interestingly, heme binding to guanine quadruplexes (G4) in DNA also has the potential to regulate gene expression ^60,61^. We have discovered several previously unknown interactions between heme and nucleosomal histones H2B and H4, as well as nucleosome remodeling proteins in response to heme depletion (**Figure 6**). Taken together, with emerging evidence from previous work, our results suggest that heme-nucleosome interactions may participate in chromatin remodeling and gene expression.

If dysregulated, labile heme can also induce DNA damage ^24–26,62^, consistent with its well-known cytotoxic effects. Heme has also been shown to interact with DNA protective or DNA repair proteins. Heme binding by DNA-protective proteins from starved cells (known as *DPs*) has been shown to protect DNA from damage in *Porphyromonas gingivalis* ^63^. In mammals, excess cytotoxic heme is catabolized by the heme oxygenase, HO-1, which also modulates ATM-dependent DNA repair ^64^. Moreover, mice lacking HO-1 undergo dysregulated expression of histone γ-H2A and an increase in DNA damage. Consistent with these reports, we have identified connections between heme and DNA repair proteins that could suggest a direct role of heme binding in the control of genome integrity. Specifically, we found heme binding proteins associated with both nucleotide excision and base excision repair processes including Ddb1 and Parp1, and a protein kinase sensor of DNA damage, Prkdc (**Figure 6**). We have also shown here that variants of histone H2B, which bind with specificity to hemin agarose, undergo a dramatic increase in expression upon heme depletion (**Figure 2 & 3**). This observation is similar to the relationship between HO-1 and histone γ-H2A and supports the hypothesis of a functional connection between heme, nucleosomes, chromatin remodeling and DNA damage repair. Integral to this correlation, we also found several heme binding proteins associated with DNA replication including helicases (e.g. G3BP1), topoisomerases (e.g. Top2a & Top2b), and mitochondrial single stranded binding protein (Ssbp1). Thus, in addition to the remodeling of chromatin, our findings support a direct role for heme in genome integrity that includes the detection and repair of DNA damage.

### 4.2 | An emerging role for heme as a feedback signal in post-transcriptional regulation

We observed several RNA processing proteins that bind specifically to hemin agarose. For example, we discovered proteins involved in tRNA and rRNA processing, such as ThumpD1 and FBL, and in RNA stability, such as SSB (**Table S4**). However, most notable were proteins involved in mRNA splicing and packaging, including multiple snRNPs and hnRNPs (**Figure 6**). Transcriptome-wide analyses of patients with myelodysplasia, wherein aberrant splicing is common, recently revealed that heme biosynthesis is major target of aberrant splicing caused by mutations in spliceosome subunit Sf3b1 ^65^. A report published in tandem showed that the long non-coding RNA, UCA1, effectively stabilizes the mRNA of Alas1 (a key heme biosynthetic enzyme) through a direct interaction with splicing factor Hnrnp1, thereby promoting heme biosynthesis ^66^. Taken together with our data, the evidence suggests a direct role for heme in splicing, including the splicing of mRNAs encoding heme biosynthetic proteins. We speculate that these processes could be mediated through feedback signaling wherein excess labile heme modulates spliceosome function. Our findings extend the role of heme in mRNA maturation beyond its established role in regulating the microRNA processing protein, DGCR8 ^67–72^.

### 4.3 | Heme regulation of proteostasis

Heme has well-established effects on the abundance of proteins – defined by the steady-state between protein synthesis and degradation – achieved through a variety of mechanisms. In some cases, heme-protein interactions are essential for maintaining the stability of HBPs. For example, the interaction between hemoglobin and heme is essential not only for oxygen transport but also to maintain stability of the protein itself ^47^. In other cases, heme promotes protein degradation, as in the case of Alas1, wherein heme functions as a feedback signal triggering ubiquitin-mediated degradation of the protein ^48^. Heme has also been shown to interact directly with the 26S proteasome and control its activity under hyper-elevated heme stress ^73^, and functions as a stoichiometric and catalytic down-regulator of the N-end rule pathway by altering arginylation rates for oxidized cysteine, aspartate and glutamate N-terminal residues that drive degradation by N-end rule E3 ubiquitin ligases ^74^. Consistent with these observations, we identified Got1 and Got2 aspartate aminotransferases as highly specific hemin-agarose binding proteins that were also unique to the SA+ heme depleted condition (**Figure 6**). These proteins are essential for the interconversion of glutamate from aspartate or cysteine, each of which is a destabilizing residue in the N-end rule pathway.

Heme has recently been shown to bind the ribosome both *in vitro* and *in vivo* by association with rRNA G-quadruplexes ^38^. G4s form in ribosomal RNAs of birds and mammals. The interactions of these G4s with heme is currently thought to serve as a buffer for labile heme that may in turn regulate heme metabolism and trafficking. Our recent studies and work by others have confirmed this interaction through the enrichment of large and small ribosomal subunits (**Figure 6**) ^40^. These ribosomes are most likely captured indirectly through heme/G4 as opposed to direct heme/protein interactions, although several ribosomal subunits harbor putative heme regulatory motifs (HRMs) that suggest ribosomal protein/heme interactions are also possible ^40^. Indeed, a hemin-affinity based approach such as the one used here cannot distinguish between protein versus non-protein-based interactions, nor can it distinguish between direct and indirect interactions, and therefore the enrichment of functional protein complexes could be facilitated by both heme-direct and indirect interactions. Whether the prominent interactions between heme and the ribosome alters the functionality of, or confers a distinctive function for the ribosome remains unknown. However, we have also identified several putative HBPs that serve accessory roles in protein translation, such as aminoacyl tRNA synthetases such as IARS and MARS that are essential for the formation of charged isoleucyl and methionyl tRNA, among other tRNA and rRNA processing enzymes described above. Thus, we speculate that heme can not only control protein degradation but may regulate protein synthesis through a variety of possible mechanisms.

Emerging evidence for heme/RNA/protein interactions such as those within the ribosome suggests that the RNA-binding subset of the proteome may be a particular target for heme-based regulation. For example, long non-coding RNAs also adopt G4 structures ^75^, interact with proteins, and can regulate heme biosynthesis ^66^, which suggests a possible role for heme in these RNA/protein interactions. In support of this hypothesis, we found that 48% of all proteins identified in the collection of hemin agarose-specific SA+/− results (102/213 proteins) are classified as RNA binding (GO:0003723; 5.6x enrichment; Q<3.24E-47) (**Figure S3**). Most of these RNA binding proteins (69/102) are not ribosomal proteins, suggesting the possibility that heme may interface with several of the identified targets through an RNA intermediary in addition or in place of direct heme/protein interaction.

### 4.4 | Involvement of heme in post-translational processes

We discovered hemin agarose-specific interactions with proteins involved in a wide range of post-translational processes including protein/vesicle trafficking, chaperone mediated protein folding, and energy metabolism. Heme has been proposed to be biodistributed as cargo in vesicular trafficking pathways between the mitochondrion, ER, golgi, and plasma membrane ^9,37,76–78^. We found prominent and specific enrichment of cytoskeletal proteins involved in endosomal trafficking to the plasma membrane, trafficking from plasma membrane to early endosomes, as well as proteins involved in cell polarity, movement, and mitosis (**Figure 6; Table S6**).

We observed several cytoplasmic and mitochondrial heat shock proteins in our analysis (**Figure 6; Table S6**). In addition to their well-established roles in protein folding, quality control, and the stress response ^79^, HSPs are also involved in iron/sulfur cluster biogenesis, important for heme biosynthesis ^80^; and in facilitating catalytic activities including intra-protein electron transfer to heme cofactors ^81^. Of note, HSPs have also been directly implicated in heme insertion into hemoproteins. For example, Hsp90 interacts with and is required for heme insertion into hemoglobin ^82^, nitric oxide synthase ^83^, and soluble guanylate cyclase ^84^. Heme has an established role in stabilizing protein folding of HBPs ^85^. Additionally, through its cytotoxic ability to promote the formation of reactive oxygen species (ROS) and to promote Nrf2-mediated transcription of redox stress response genes, excess labile heme can promote protein misfolding and aggregation ^86^. Thus, we speculate that our detection of heme and heat shock protein interactions could be indicative of a feedback mechanism wherein excess labile heme released upon stress signal directly to cytoprotective chaperones that stabilize the proteome.

### 4.5 | Conclusions and future outlook

DAsHA is a new approach that seeks to discover novel hemoproteins by depleting endogenous HBPs of their heme to enhance their enrichment through traditional hemin agarose chromatography. By abolishing exchange labile heme, but not total heme, we sought to identify proteins that constitute the network of labile HBPs. Indeed, our approach did not identify canonical high affinity heme proteins, but rather uncovered an array of previously unknown HBPs, thereby expanding the landscape of the hemoproteome. While we identified GAPDH in our studies, a protein previously implicated in heme trafficking, we did not identify known heme transporters, HRG1^87^, FLVCR1^88^, and FLVCR2 ^89^, or heme trafficking factors like PGRMC1 and PGRMC2 ^90^. FLVCR2 and HRG1 were not identified since peptides from these proteins were not detected at the whole proteome level. On the other hand, peptides from PGRMC1/2 and FLVCR1 were detected at the whole proteome-level, but not found associated with hemin-agarose, possibly because the heme propionates, which are used in the conjugation of heme to the hemin-agarose resin, are unavailable for interactions with proteins that require such engagement ^91,92^, as is the case for PGRMC1/2 ^93^. It remains to be determined if a subset of the HBPs identified in this study play roles in heme homeostasis or are novel heme-dependent or regulated factors. The identification of putative signatures of heme regulatory motifs ^3,94–98^, and the development of computational tools to predict such heme binding sites ^99^, will guide our future work to establish functional consequences of heme binding to the HBPs identified herein.

Our studies are complemented by Tsolaki et al. ^40^, in which heme interacting proteins were identified from cells in the presence versus absence of excess hemin (as opposed to our work in which heme is depleted below physiological conditions). Overlap between the two datasets provides a view of similarities and differences that not only underscore reproducibility of shared findings, but also highlight unique findings that were achieved only through heme depletion. Shared findings between the two studies include heat shock proteins, cytoskeletal proteins, ATP synthases, heterogenous ribonucleoproteins, proteasomal subunits, splicing factors and the DNA damage response protein Ddb1. Several unique findings from our work therefore include proteins involved in vesicular trafficking, nucleosome and nucleosome remodeling, the spliceosome, and DNA replication (**Figure 6**). In many of these cases we observed enrichment only upon heme depletion, supporting the conclusion that DAsHA can provide a unique view into the heme binding subset of the human proteome and can complement other proteomics approaches.

## 5| ACKNOWLEDGEMENTS

We acknowledge support from US National Institutes of Health (NIH) grants GM118744 (to A.R.R. and M.P.T.), GM145350 (to A.R.R.), ES025661 (to A.R.R.), NS123168 (to A.R.R.), a US National Science Foundation (NSF) grant 1552791 (to A. R. R.), and the Georgia Institute of Technology Blanchard and Vasser-Woolley fellowships (to A. R. R.).

## Notes

### Competing Interest Statement

The authors have declared no competing interest.

