## Supplemental Information for "Depletion Assisted Hemin Affinity (DAsHA) Proteomics Reveals an Expanded Landscape of Heme Binding Proteins"

Contains Figures S1 – S5

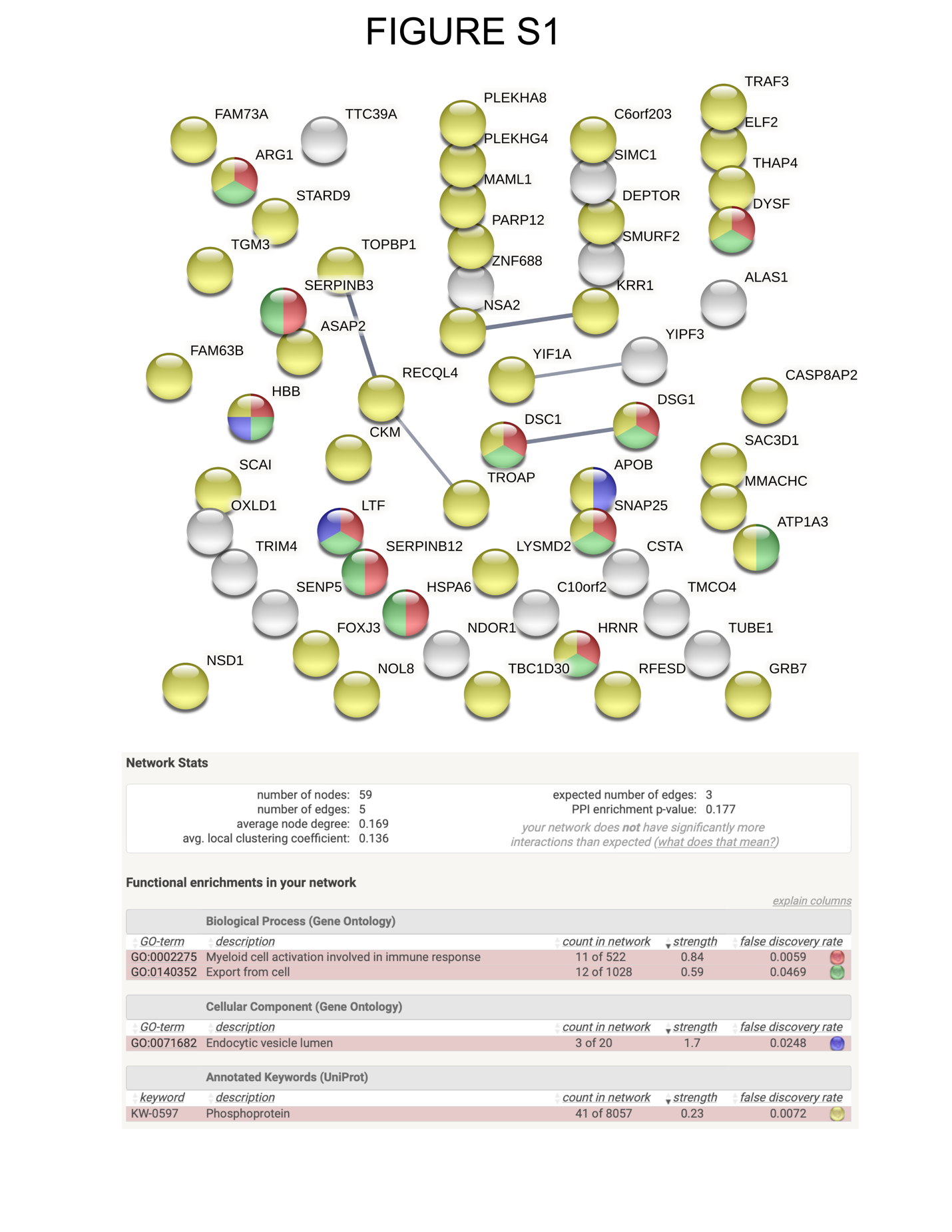

**Figures S1.** Proteins undergoing extreme abundance changes in response to heme depletion with succinylacetone. Only High confidence physical interactions are shown (grey lines).

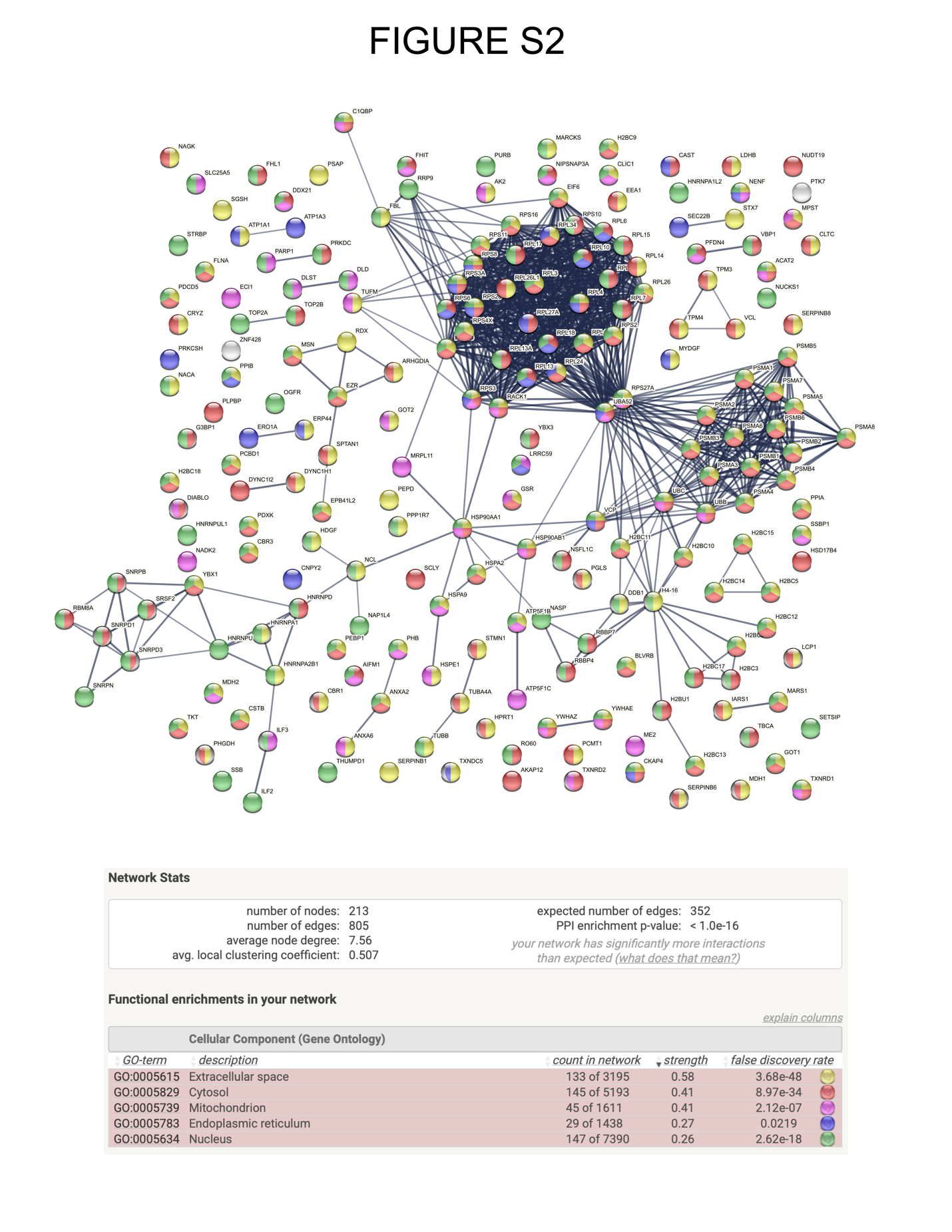

**Figure S2.** **The heme-binding proteins identified in this study localize to all major locations in the cell.** Full STRING network for proteins classified as moderate or high specificity hemin agarose binding in either SA- or SA+ conditions. Proteins are clustered by MCL method (default). Only high confidence physical interactions are shown (grey lines). Color coding key is show in table at bottom.

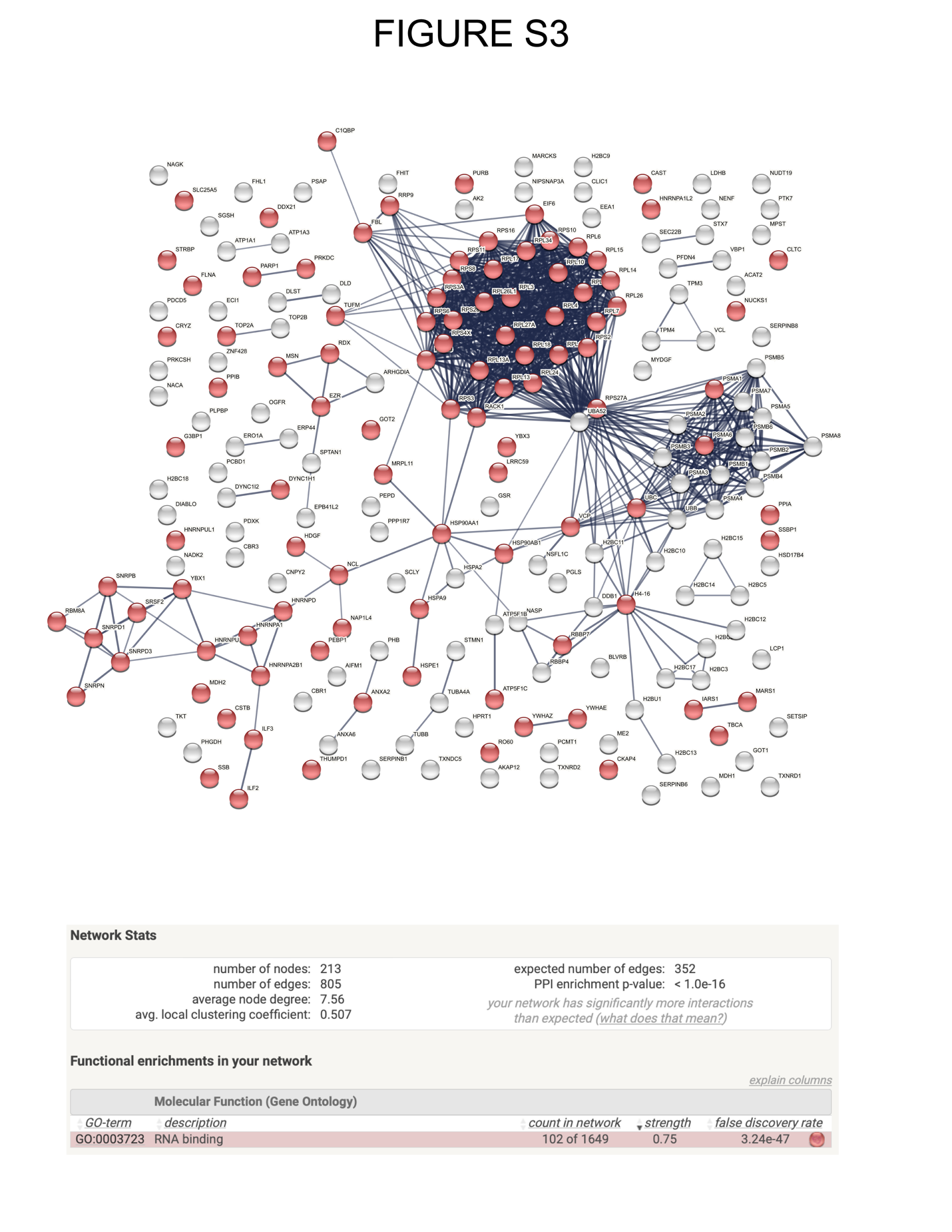

**Figure S3. Nearly 50% of all hemin agarose specific proteins are RNA binding proteins.** Full STRING network for proteins classified as moderate or high specificity hemin agarose binding in either SA- or SA+ conditions. Proteins are clustered by MCL method (default). Only high confidence physical interactions are shown (grey lines). Color coding key is show in table at bottom.

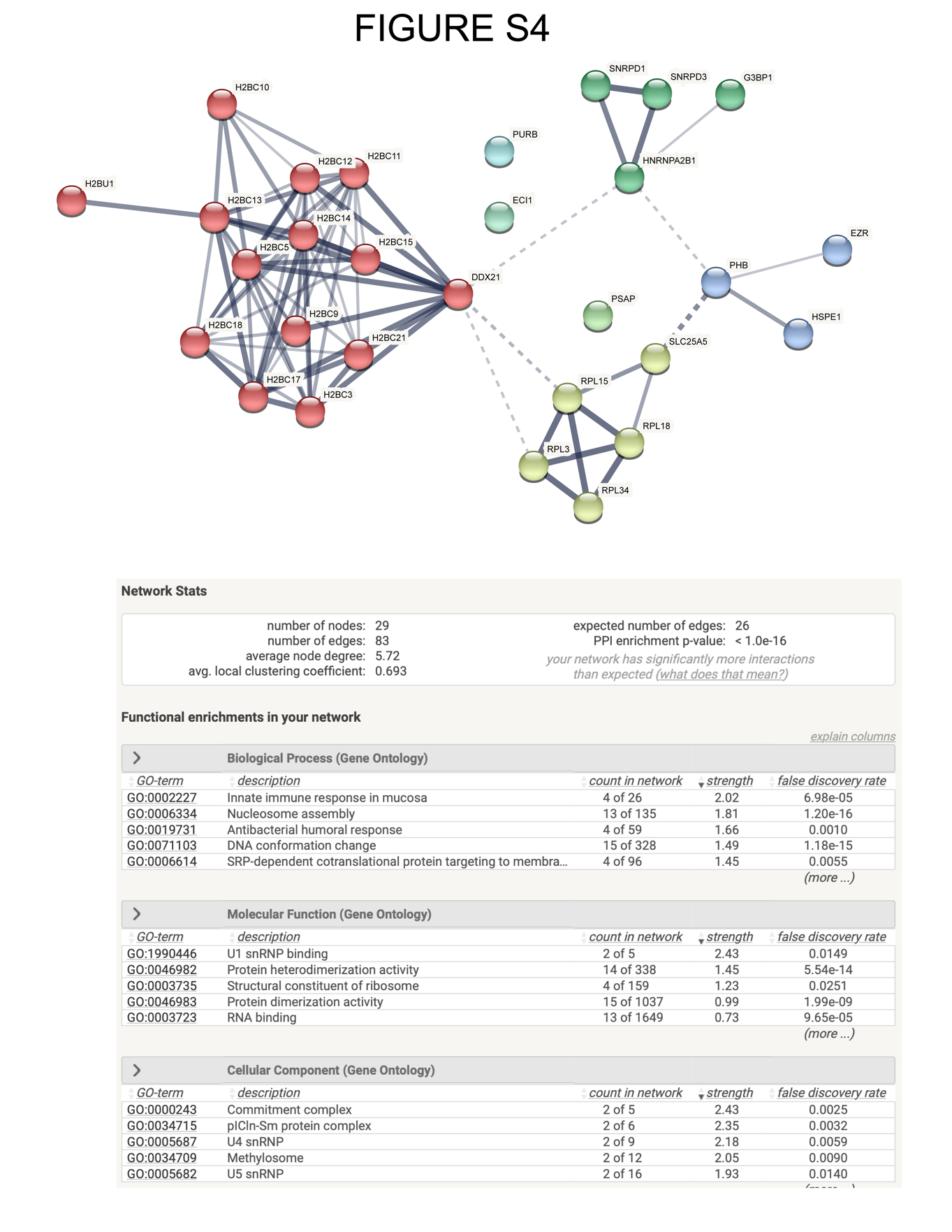

Figure S4. Subset of hemin agarose-interacting proteins that also undergo extreme abundance change in response to heme depletion with succinylacetone. Proteins shown are those with large change in abundance after SA treatment (Log_2_ SA+/SA- >5) that also undergo significant enrichment with hemin agarose compared to sepharose. Bottom table shows major GO terms for this network.

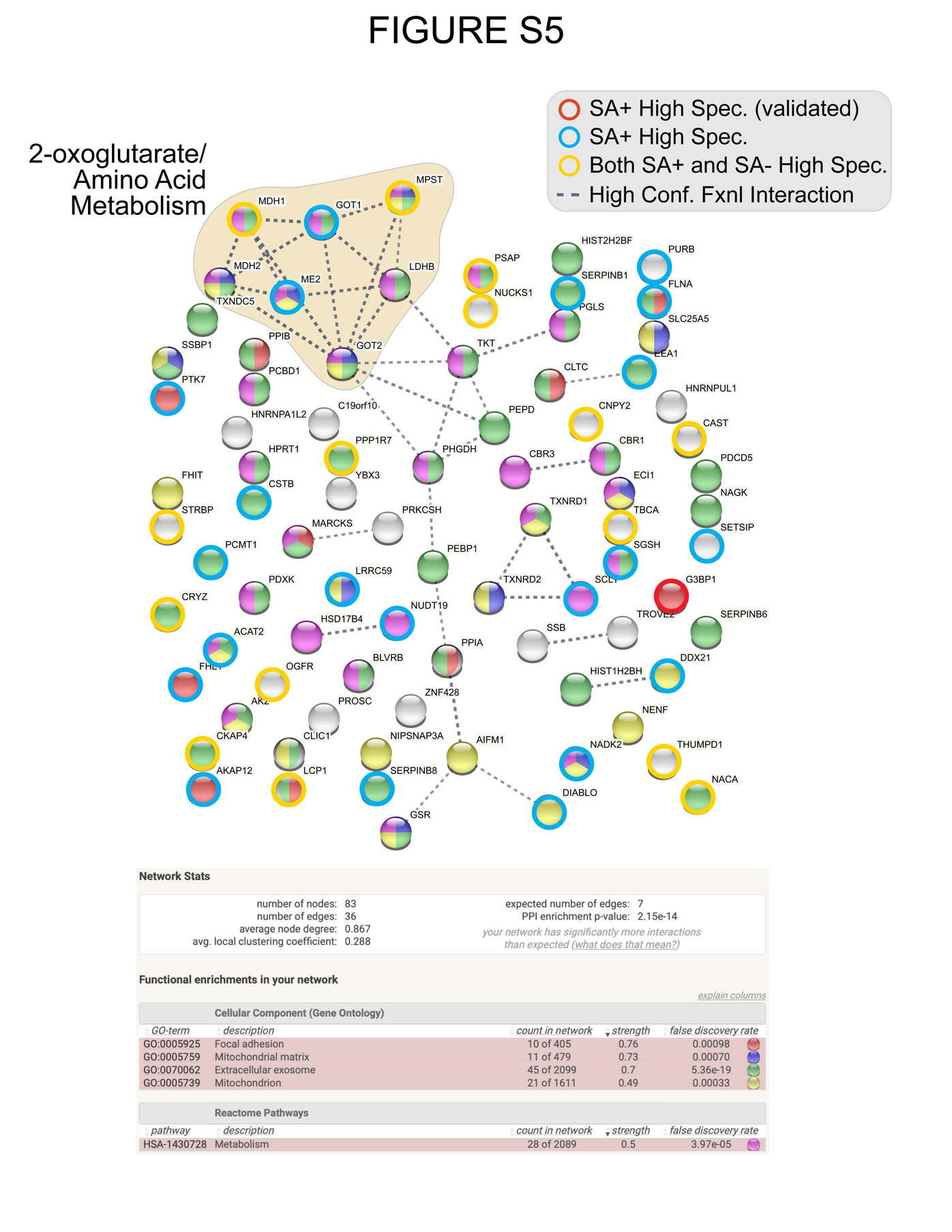

**Figure S5. Subset of hemin agarose-interacting proteins that do not exhibit high confidence physical interactions.** Proteins were clustered using the MCL inflation method in STRING. Spheres encircled with red, blue, or yellow rings were detected in the high specificity class (exclusively detected from hemin agarose but not sepharose enrichment) while spheres without rings were detected in the moderate specificity class (2 to 70-fold hemin agarose versus sepharose enrichment). Amino acid metabolism cluster is also shown in Figure 6.
